# Brain Water as a Function of Age and Weight in Normal Rats

**DOI:** 10.1101/2021.03.18.436025

**Authors:** Allan Gottschalk, Susanna Scafidi, Thomas J. K. Toung

## Abstract

Rats are frequently used to study water content of normal and injured brain, as well as changes in response to various osmotherapeutic regimens. Magnetic resonance imaging in humans has shown that brain water content declines with age as a result of progressive myelination. The purpose of this study was to quantify changes in brain water content during rat development and aging. Brain water content was measured by standard techniques in 129 normal male Sprague-Dawley rats that ranged in age (weight) from 13 to 149 days (18 to 759 g). Overall, the results demonstrated a decrease from 85.59% to 76.56% water content with increasing age (weight). Nonlinear allometric functions relating brain water to age and weight were determined. These findings provide age-related context for prior rat studies of brain water, emphasize the importance of using similarly aged controls in studies of brain water, and indicate that age-related changes in brain water content are not specific to humans.

## 1. Introduction

Rats continue to be used extensively in laboratory medicine and have been the source of considerable animal data regarding water content of the normal and injured brain (1–3). Many of these studies have used rats to examine the extent to which various osmotherapeutic regimens can reduce brain water content (2, 3). Magnetic resonance imaging has shown that brain water content decreases with increasing gestational age in humans and rabbits (4, 5). This decline in water content reflects increasing myelination (6, 7). Such findings suggest that, at least in humans and rabbits, not only does brain water vary with age, but that brain water content is tightly coupled to age. The purpose of this study was to determine whether brain water content in normal rats is also dependent on age and to thereby provide important age-related context for prior studies of brain water content.

## 2. Materials and Methods

All animal protocols were approved by the Johns Hopkins Institutional Animal Care and Use Committee and were carried out in male Sprague-Dawley rats (Charles River Laboratories, Inc., Wilmington, DE, USA). Spontaneously breathing rats were initially anesthetized by isoflurane (1-2%) in an oxygen-air mixture delivered through a face mask. Then, their weight was determined, and the animals were sacrificed under deep anesthesia. The whole brain was procured, weighed, and dried at 35-38°C for 2 days. Brain water content was calculated as follows: % H_2_O = (1 – dry weight/wet weight) × 100 (8). A total of 129 animals were studied in 12 groups. Each group was of a different average age and weight when ordered from the supplier, and ranged in number from 2 to 20 animals. The median [IQR] group size was 9 [8, 12] rats.

### 2.1 Statistical Analysis

All data are presented as median [IQR], with linear and nonlinear regression used to establish continuous relationships between brain water content, age, and weight. The linear regression was checked to determine if additional terms would contribute to the quality of the fit. The mean and standard error of the mean (SE) are reported for each regression parameter. The nonlinear functions to which we fit the data were initially chosen more for convenience than to validate any preconceived theoretical concept regarding the nature of these relationships but, as detailed in the Discussion, relate to classical concepts of allometric scaling (9). Analysis was facilitated with Mathematica 12.1 (Wolfram Research, Inc., Champaign, IL, U.S.A.).

## 3. Results

Overall, the 129 male Sprague-Dawley rats ranged in age (weight) from 13 to 149 days (18 to 759 g). Brain water content decreased from 85.59% to 76.56% with increasing age (weight). Results from the entire study population are shown in Fig. 1, with associated detail in Table 1. Fig. 1A illustrates a monotonic curvilinear decrease in brain water content with weight that begins to flatten for the largest animals. The variation observed in brain water between the extremes of weight was almost 10%. Because weight and age for the study animals was linearly related (Fig. 1B), in Fig. 1C, brain water as a function of age was fit with the same function as used in Fig. 1A.

**Figure 1.**
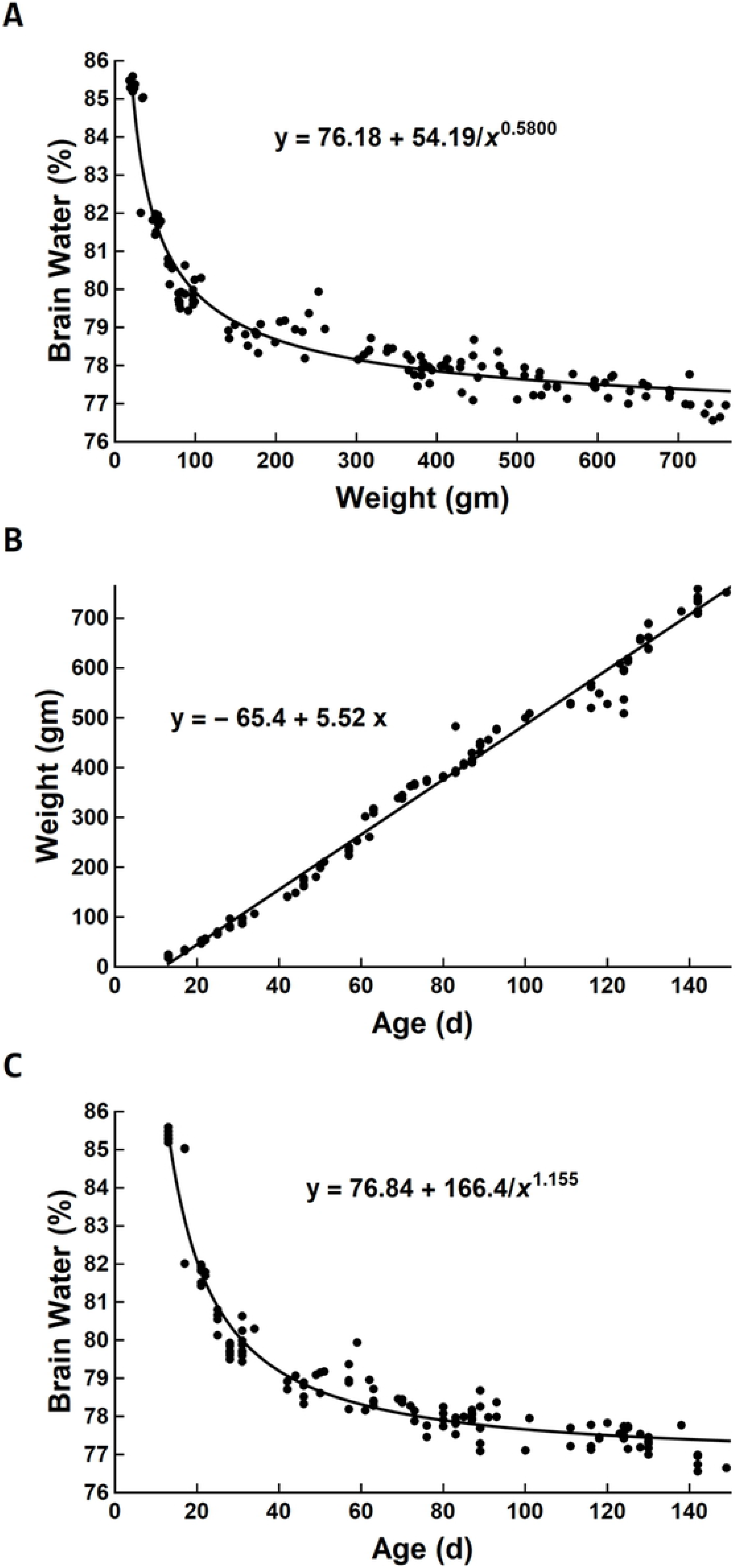
Nonlinear relationship between brain water content and weight (A), linear relationship between weight and age (B), and nonlinear relationship between brain water content and age (C) for all 129 rats from the study. The corresponding regression equations are given in each panel, where the functional forms of the nonlinear relationships in A and C are the same but with different parameters. A quadratic term for the linear regression in panel B was not contributory (*p* = 0.30). All parameters of the regression equations in each panel are significantly different from zero (*p* < 0.001) and are given along with their standard errors in Table 1.

**Table 1.** Parameters for regression equations for panels of Fig. 1. All parameters are significantly different from zero (*p* < 0.001).

| **Regression**<br>**equation** | **Parameter 1**<br>**mean (SE)** | **Parameter 2**<br>**mean (SE)** | **Parameter 3**<br>**mean (SE)** | **R<sup>2</sup>** |
| --- | --- | --- | --- | --- |
| Fig. 1, panel A | 76.18 (0.26) | 54.19 (1.18) | 0.5800 (0.0431) | >0.99 |
| Fig. 1, panel B | -65.4 (4.2) | 5.52 (0.05) | NA | 0.99 |
| Fig. 1, panel C | 76.84 (0.16) | 166.42 (1.61) | 1.155 (0.066) | >0.99 |

NA = not applicable; SE, standard error of the mean.

## 4. Discussion

For rats ranging in age (weight) from 13 to 149 days (18 to 759 g), brain water content declined by almost 10% of total brain weight. This decrease should be compared to studies of osmotherapy intended to determine the reduction in brain water associated with such therapy, where typical changes in brain water are on the order of 1.5% (2, 3). This finding emphasizes the importance of using control animals of identical age in experiments involving measurement of brain water, particularly when studying animals at young ages, when the brain water-age curve is particularly steep. The magnitude of the decrease in brain water content with age and its curvilinear shape with a steep initial descent is similar to what has been observed in humans and rabbits by magnetic resonance imaging (4, 5). The data from magnetic resonance imaging reflect progressive myelination with development (6, 7) and are also consistent with the age-dependent expression of aquaporin channels in the brain (10).

The nonlinear equations used to fit the data were initially chosen empirically for their utility in doing just that. However, we retained them, not only because of their ability to fit the data but because the nonlinear term of the form ax^b^ in the equation used to fit the data, where a and b are coefficients and x is the independent variable, is frequently utilized in allometric models (9). The clear linear relationship between weight and age for the available data permitted brain water content to be related to each of these variables using equations of the same functional form. A much smaller study that used Wistar rats over approximately the same age range as those in our study (11) reported an inflection point in the weight-age curve that was clearly not observed here. Whether the lack of a similar inflection point in the current somewhat larger study represents differences in how the animals were fed and maintained, differences in the type of rat (Wistar vs. Sprague-Dawley), or something more fundamental is not apparent.

One additional limitation of our study was the use of only male rats. This single-sex study is consistent with prior studies of osmotherapy with mannitol (2, 3), in which researchers used only male animals to avoid any possible confounding due to the established vascular effects of estrogen, particularly altered membrane permeability (12).

## 5. Conclusions

In normal male Sprague-Dawley rats, brain water content expressed as a percentage of total brain weight declines in a steep curvilinear fashion by almost 10% over the early phase of life and asymptotes in adulthood.

## Acknowledgements

The authors are grateful for the editorial assistance of Claire Levine, MS, ELS, Manager, Editorial Services, Department of Anesthesiology and Critical Care Medicine, Johns Hopkins University, Baltimore, MD, USA.

